# SARS-coronavirus-2 replication in Vero E6 cells: replication kinetics, rapid adaptation and cytopathology

**DOI:** 10.1101/2020.04.20.049924

**Authors:** Natacha S. Ogando, Tim J. Dalebout, Jessika C. Zevenhoven-Dobbe, Ronald W. Limpens, Yvonne van der Meer, Leon Caly, Julian Druce, Jutte J. C. de Vries, Marjolein Kikkert, Montserrat Bárcena, Igor Sidorov, Eric J. Snijder

## Abstract

The sudden emergence of severe acute respiratory syndrome coronavirus 2 (SARS-CoV-2) at the end of 2019 from the Chinese province of Hubei and its subsequent pandemic spread highlight the importance of understanding the full molecular details of coronavirus infection and pathogenesis. Here, we compared a variety of replication features of SARS-CoV-2 and SARS-CoV and analysed the cytopathology caused by the two closely related viruses in the commonly used Vero E6 cell line. Compared to SARS-CoV, SARS-CoV-2 generated higher levels of intracellular viral RNA, but strikingly about 50-fold less infectious viral progeny was recovered from the culture medium. Immunofluorescence microscopy of SARS-CoV-2-infected cells established extensive cross-reactivity of antisera previously raised against a variety of nonstructural proteins, membrane and nucleocapsid protein of SARS-CoV. Electron microscopy revealed that the ultrastructural changes induced by the two SARS viruses are very similar and occur within comparable time frames after infection. Furthermore, we determined that the sensitivity of the two viruses to three established inhibitors of coronavirus replication (Remdesivir, Alisporivir and chloroquine) is very similar, but that SARS-CoV-2 infection was substantially more sensitive to pre-treatment of cells with pegylated interferon alpha. An important difference between the two viruses is the fact that - upon passaging in Vero E6 cells - SARS-CoV-2 apparently is under strong selection pressure to acquire adaptive mutations in its spike protein gene. These mutations change or delete a putative ‘furin-like cleavage site’ in the region connecting the S1 and S2 domains and result in a very prominent phenotypic change in plaque assays.

## INTRODUCTION

For the first time in a century, societies and economies worldwide have come to a near-complete standstill due to a pandemic outbreak of a single RNA virus. This virus, the severe acute respiratory syndrome coronavirus 2 (SARS-CoV-2) (1) belongs to the coronavirus (CoV) family, which is thought to have given rise to zoonotic introductions on multiple previous occasions during the past centuries. Coronaviruses are abundantly present in mammalian reservoir species, including bats (2), and should now be recognized definitively as a continuous zoonotic threat with the ability to cause severe human disease and explosive pandemic transmission.

To date, seven CoVs that can infect humans have been identified, which segregate into two classes. On the one hand, there are four endemic human CoVs (HCoVs), the first of which were identified in the 1960’s, annually causing a substantial number of common colds (3, 4). On the other hand, we now know of (at least) three zoonotic CoVs that have caused outbreaks in the human population recently: severe acute respiratory syndrome coronavirus (SARS-CoV) (5, 6) in 2002-2003, Middle East respiratory syndrome-coronavirus (MERS-CoV) (7, 8) since 2012 (and probably earlier) and the currently pandemic SARS-CoV-2 (9, 10). The latter agent emerged near Wuhan (People’s Republic of China) in the fall of 2019 and its animal source is currently under investigation (11-13). Transmission to humans of SARS-CoV and MERS-CoV was attributed to civet cats (14) and dromedary camels (15), respectively, although both species may have served merely as an intermediate host due to their close contact with humans. All three zoonotic CoVs belong to the genus *betacoronavirus* (beta-CoV), which is abundantly represented among the CoVs that circulate in the many bat species on this planet (2, 16-19). The genetic diversity of bat CoVs and their phylogenetic relationships with the four known endemic HCoVs (OC43, HKU1, 229E and NL63; the latter two being alpha-CoVs) suggests that also these may have their evolutionary origins in bat hosts, for most of them probably centuries ago (20).The potential of multiple CoVs from different genera to cross-species barriers had been predicted and documented previously (2, 16-19, 21, 22), but regrettably was not taken seriously enough to invest more extensively in prophylactic and therapeutic solutions that could have contributed to rapidly containing an outbreak of the current magnitude.

Compared to other RNA viruses, CoVs possess an unusually large positive-sense RNA genome with a size ranging from 26 to 34 kilobases (23). The CoV genome is single-stranded and its 5’-proximal two-thirds encode for the large and partially overlapping replicase polyproteins pp1a and pp1ab (4,000-4,500 and 6,700-7,200 amino acids long, respectively), with the latter being a C-terminally extended version of the former that results from ribosomal frameshifting. The replicase polyproteins are processed into 16 cleavage products (non-structural proteins, nsps) by two internal proteases, the papain-like protease (PL^pro^) in nsp3 and the 3C-like or ‘main’ protease (M^pro^) in nsp5 (24). Specific trans-membrane nsps (nsp3, 4 and 6) than cooperate to transform intracellular membranes into a viral replication organelle (RO) (25) that serves to organize and execute CoV RNA synthesis, which entails genome replication and the synthesis of an extensive nested set of subgenomic (sg) mRNAs. The latter are used to express the genes present in the 3’-proximal third of the genome, which encode the four common CoV structural proteins (spike (S), envelope (E), membrane (M) and nucleocapsid (N) protein) and the ‘so-called’ accessory protein genes, most of which are thought to be involved in the modulation of host responses to CoV infection (26). The CoV proteome includes a variety of potential targets for drug repurposing or *de novo* development of specific inhibitors of e.g. viral entry (S protein) or RNA synthesis (27). The latter process depends on a set of enzymatic activities (24) including an RNA-dependent RNA polymerase (RdRp; in nsp12), RNA helicase (in nsp13), two methyltransferases involved in mRNA capping (a guanine-N7-methyltranferase in nsp14 and a nucleoside-2’-O-methyltransferase in nsp16) and a unique exoribonuclease (ExoN, in nsp14) that promotes the fidelity of the replication of the large CoV genome (28). Other potential drug targets are the transmembrane proteins that direct the formation of the viral RO, several less well characterised enzymatic activities and a set of smaller nsps (nsp7-10) that mainly appear to serve as cofactors/modulators of other nsps.

The newly emerged SARS-CoV-2 was rapidly identified as a CoV that is relatively closely related to the 2003 SARS-CoV (9, 29, 30).The two genome sequences are about ∼80% identical and the organization of open reading frames is essentially the same. The overall level of amino acid sequence identity of viral proteins ranges from about 65% in the least conserved parts of the S protein to about 95% in the most conserved replicative enzyme domains, prompting the coronavirus study group of the International Committee on the Taxonomy of Viruses to classify the new agent within the species *Severe acute respiratory syndrome-related coronavirus*, which also includes the 2003 SARS-CoV (1). The close phylogenetic relationship also implies that much of our knowledge of SARS-CoV molecular biology, accumulated over the past 17 years, can probably be translated to SARS-CoV-2. Many reports posted over the past months have described such similarities, including the common affinity of the two viruses for the angiotensin-converting enzyme 2 (ACE2) receptor (9, 31). This receptor is abundantly expressed in Vero cells (African green monkey kidney cells). Since 2003, Vero cells have been used extensively for SARS-CoV research in cell culture-based infection models by many laboratories, including our own.

We set out to establish the basic features of SARS-CoV-2 replication in Vero cells and compare it to the Frankfurt-1 SARS-CoV isolate from 2003 (32, 33). When requesting virus isolates (February 2020), and in spite of the rapidly emerging public health crisis, we were confronted - not for the first time - with administrative hurdles and discussions regarding the alleged ‘ownership’ of virus isolates cultured from (anonymous) clinical samples. From a biological and evolutionary point of view, this would seem a strangely anthropocentric consideration, but it ultimately forced us to reach out across the globe to Australian colleagues in Melbourne. After checking our credentials and completing a basic material transfer agreement, they provided us (within one week) with their first SARS-CoV-2 isolate (originally named 2019-nCoV/Victoria/1/2020 and subsequently renamed BetaCoV/Australia/VIC01/2020; (34), which will be used throughout this study. Until now, this isolate has been provided to 17 other laboratories worldwide to promote the rapid characterization of SARS-CoV-2, in this critical time of lockdowns and other preventive measures to avoid a collapse of public health systems.

In this report, we describe a comparative study of the basic replication features of SARS-CoV and SARS-CoV-2 in Vero E6 cells, including growth kinetics, virus titres, plaque phenotype and an analysis of intracellular viral RNA and protein synthesis. Additionally, we analysed infected cells by light and electron microscopy, and demonstrated cross-reactivity of 13 available SARS-CoV-specific antisera (recognising 10 different viral proteins) with their SARS-CoV-2 counterparts. Finally, we established the conditions for a medium-throughput assay to evaluate basic antiviral activity and assessed the impact of some known CoV inhibitors on SARS-CoV-2 replication. In addition to many anticipated similarities, our results also established some remarkable differences between the two viruses that warrant further investigation. One of them is the rapid evolution - during virus passaging in Vero cells - of a specific region of the SARS-CoV-2 S protein that contains the so-called ‘furin-like cleavage site’.

## METHODS

### Cell and virus culture

Vero E6 cells and HuH7 cells were grown as described previously (35). SARS-CoV-2 isolate Australia/VIC01/2020 (GeneBank ID: MT007544.1; (34)) was derived from a positively-testing nasopharyngeal swab in Melbourne, Australia, and was propagated twice in Vero/hSLAM cells, before being shared with other laboratories. In Leiden, the virus was passaged two more times at low multiplicity of infection (m.o.i.) in Vero E6 cells to obtain a working stock (p2 stock). SARS-CoV isolate Frankfurt 1 (36) was used to compare growth kinetics and other features with SARS-CoV-2. Infection of Vero E6 cells was carried out in phosphate-buffered saline (PBS) containing 50 µg/ml DEAE-dextran and 2% fetal calf serum (FCS; Bodinco). The inoculum was added to the cells for 1 h at 37°C, after which cells were washed twice with PBS and maintained in Eagle’s minimal essential medium (EMEM; Lonza) with 2% FCS, 2mM L-glutamine (PAA) and antibiotics (Sigma). Viral titres were determined by plaque assay in Vero E6 cells as described previously (37). For plaque picking, plaque assays were performed using our p1 stock, while using an overlay containing 1% of agarose instead of Avicel (RC-581; FMC Biopolymer). Following neutral red staining, small and large plaques were picked and used to inoculate a 9.6-cm^2^ dish of Vero E6 cells containing 2 ml of EMEM-2%FCS medium, yielding p1 virus. After 48 h, 200 µl of the culture supernatant was used to infect the next dish of cells (p2), a step that was repeated one more time to obtain p3 virus. All work with live SARS-CoV and SARS-CoV-2 was performed in biosafety laboratory level 3 facilities at Leiden University Medical Center, the Netherlands.

### Analysis of intracellular viral RNA and protein synthesis

Isolation of intracellular RNA was performed by lysing infected cell monolayers with TriPure isolation reagent (Roche Applied Science) according to the manufacturer’s instructions. After purification and ethanol precipitation, intracellular RNA samples were loaded onto a 1.5% agarose gel containing 2.2 M formaldehyde, which was run overnight at low voltage overnight in MOPS buffer (10 mM MOPS (sodium salt) (pH 7), 5 mM sodium acetate, 1 mM EDTA). Dried agarose gels were used for direct detection of viral mRNAs by hybridization with a ^32^P-labeled oligonucleotide probe (5’-CACATGGGGATAGCACTAC-3’) that is complementary to a fully conserved sequence located 30 nucleotides upstream of the 3’ end of the genome and all subgenomic mRNAs produced by SARS-CoV-2 and SARS-CoV. After hybridization, RNA bands were visualised and quantified by phosphorimaging using a Typhoon-9410 variable mode scanner (GE Healthcare) and ImageQuant TL software (GE Healthcare). In order to verify the amount of RNA loaded, a second hybridization was performed using a ^32^P-labeled oligonucleotide probe recognizing 18S ribosomal RNA (5’-GATCCGAGGGCCTCACTAAAC-3’). Protein lysates were obtained by lysing infected cell monolayers in 4x Laemmli sample buffer and were analysed by semi-dry Western blotting onto Hybond 0.2µM polyvinylidene difluoride (PVDF) membrane (GE Healthcare). Membranes were incubated with rabbit antisera diluted in PBS with 0.05% Tween-20 containing 5% dry milk (Campina). Primary antibodies were detected with a horseradish peroxidase-conjugated swine anti-rabbit IgG antibody (Dako) and protein bands were visualised using Clarity Western Blot substrate (Biorad) and detected using an Advanced Q9 Alliance imager (Uvitec Cambridge).

### Next-generation sequencing and bioinformatics analysis

SARS-CoV-2 genomic RNA was isolated from cell culture supernatants using TriPure isolation reagent (Roche Applied Science) and purified according to manufacturer’s instructions. The total amount of RNA in samples was measured using a Qubit fluorometer and RNA High Sensitivity kit (Thermo Fisher Scientific). For next-generation sequencing (NGS) library preparation, RNA (25-100 ng) was mixed with random oligonucleotide primers using the NEBNext® First Strand Synthesis Module kit for Illumina® (NEB) and incubated for 10 min at 94°C. NGS of samples was performed by a commercial service provider (GenomeScan, Leiden, the Netherlands) while including appropriate quality controls after each step of the procedure. Sequencing was performed using a NovaSeq 6000 Sequencing System (Illumina). Subsequently, sequencing reads were screened for the presence of human (GRCh37.75), mouse (GRCm38.p4), E. coli MG1655 (EMBL U00096.2), phiX (RefSeq NC_001422.1) and common vector sequences (UniVec and ChlSab1.1). Prior to alignment, reads were trimmed to remove adapter sequences and filtered for sequence quality. The remaining reads were mapped to the SARS-CoV-2 GenBank reference sequence (NC_045512.2; (38)). Data analysis was performed using Bowtie 2 (39). Raw NGS data sets for each virus sample analysed in this study are deposited in NCBI Bioproject and available under the following links: ---. Only SARS-CoV-2-specific reads were included in these data files.

To study evolution/adaptation of the S protein gene, we performed an in-depth analysis of reads covering the S1/S2 region of the S protein gene. This was done for the p2 stock and for the four virus samples of the plaque picking experiment shown in Fig. 1a. First, all reads spanning nt 23,576 to 23,665 of the SARS-CoV genome were selected. Next, reads constituting less than 1% of the total number of selected reads were excluded for further analysis. The remaining number of reads were 3,860 (p2 stock), 1,924 (S5p1), 2,263 (S5p2), 4,049 (S5p3) and 3,323 (L8p1). These reads were translated in the S protein open reading frame and the resulting amino acid sequences were aligned, grouped on the basis of containing the same mutations/deletions in the S1/S2 region and ranked by frequency of occurrence (Fig. 1b).

**Fig. 1.**
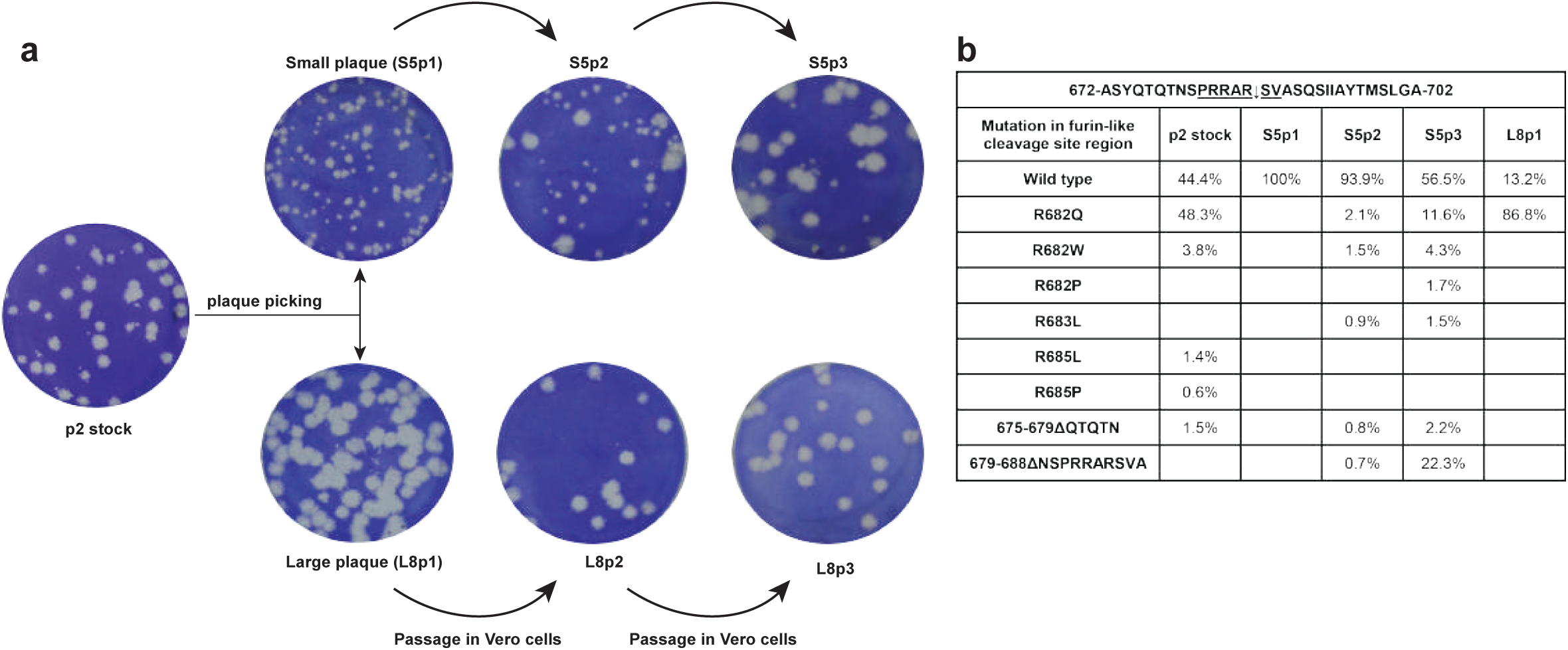
Rapid evolution of SARS-CoV-2 during passaging in Vero E6 cells. (a) Outline of a plaque picking experiment that was initiated when the p2 stock of SARS-CoV-2 Australia/VIC01/2020 showed remarkable plaque heterogeneity on Vero E6 cells (leftmost well). Following a plaque assay of the p1 virus stock, small and large plaques were picked and these virus clones were passaged three times in Vero E6 cells, while their plaque phenotype was monitored. In contrast to the large plaque viruses (example L8; bottom row), the plaque phenotype of the small plaque viruses (example S5; top row) rapidly evolved within these 3 passages. (b) Evolution/adaptation of the S protein gene during Vero E6 passaging. Overview of NGS data obtained for the p2 stock, S5p1/p2/p3 and S8p1 in the S1/S2 region of the SARS-CoV-2 S protein gene that encodes the so-called ‘furin-like cleavage site. The analysis was based on NGS reads spanning nt 23,576 to 23,665 of the SARS-CoV genome (see Methods for details) and their translation in the S protein open reading frame. Deletions are indicated with Δ followed by the affected amino acid residues.

### Antisera and immunofluorescence microscopy

The SARS-CoV-specific rabbit or mouse antisera/antibodies used in this study are listed in Table 1. Most antisera were described previously (see references in Table 1), with the exception of three rabbit antisera recognizing SARS-CoV nsps 8, 9 and 15. These were raised using full-length (His)_6_-tagged bacterial expression products (nsp8 and nsp15) or a synthetic peptide (nsp9, aa 4209-4230 of SARS-CoV pp1a), which were used to immunize New Zealand white rabbits as described previously (40, 41). Cross-reactivity of antisera to SARS-CoV-2 targets was evaluated microscopically by immunofluorescence assay (IFA) and for some antisera (nsp3 and N protein) also by Western blot analysis. Double-stranded RNA was detected using mouse monoclonal antibody J2 from Scicons (42).

**Table 1.** SARS-CoV-specific antisera used and their cross-reactivity with corresponding SARS-CoV-2 targets.

| SARS-CoV antiserum | function of target | antigen type | antibody type | IFA signal* | reference |
| --- | --- | --- | --- | --- | --- |
| nsp3 (DGD7) | transmembrane replicase protein, containing PL<sup>pro</sup> | bacterial expression product | rabbit polyclonal | ++ | (47) |
| nsp4 (FGQ4) | transmembrane replicase protein | synthetic peptide | rabbit polyclonal | ++ | (109) |
| nsp5 (DUE5) | M<sup>pro</sup> | bacterial expression product | rabbit polyclonal | + | (47) |
| nsp6 (GBZ7) | transmembrane replicase protein | synthetic peptide | rabbit polyclonal | - | (109) |
| nsp8 (DUK4) | RNA polymerase co-factor | bacterial expression product | rabbit polyclonal | ++ | (47) |
| nsp8 (39-12) | RNA polymerase co-factor | bacterial expression product | mouse monoclonal | ++ | unpublished |
| nsp9 (HLJ5) | RNA-binding protein | synthetic peptide | rabbit polyclonal | ++ | unpublished |
| nsp13 (CQS2) | RNA helicase | synthetic peptide | rabbit polyclonal | ++ | (47) |
| nsp15 (HLT5) | endoribonuclease | bacterial expression product | rabbit polyclonal | + | unpublished |
| nsp15 (BGU6) | endoribonuclease | synthetic peptide | rabbit polyclonal | + | (47) |
| M (EKU9) | membrane protein | synthetic peptide | rabbit polyclonal | + | (47) |
| N (JUC3) | nucleocapsid protein | bacterial expression product | rabbit polyclonal | + | (35) |
| N (46-4) | nucleocapsid protein | bacterial expression product | mouse monoclonal | ++ | (41) |

\* ++, strongly positive; +, positive; -, negative.

Cells were grown on glass coverslips and infected as described above (43). At 12, 24, 48 or 72 h p.i., cells were fixed overnight at 4°C using 3% paraformaldehyde in PBS (pH 7.4). Cells were washed with PBS containing 10 mM glycine and permeabilized with 0.1% Triton X-100 in PBS. Cells were incubated with antisera diluted in PBS containing 5% FCS. Secondary antibodies used were an Alexa488-conjugated goat anti-rabbit IgG antibody (Invitrogen), a Cy3-conjugated donkey anti-rabbit IgG antibody (Jackson ImmunoResearch Laboratories) and an Alexa488-conjugated goat anti-mouse IgG antibody (Invitrogen). Nuclei were stained with 1 µg/ml Hoechst 33258 (ThermoFischer). Samples were embedded using Prolong Gold (Life Technologies) and analysed with a Leica DM6B fluorescence microscope using LASX software.

### Electron microscopy

Vero E6 cells were grown on TC treated Cell Star dishes (Greiner Bio-One) and infected at an m.o.i. of 3, or mock-infected. Cells were fixed after 6, 8 and 10 h p.i. for 30 min at room temperature with freshly prepared 2% (vol/vol) glutaraldehyde in 0.1 M cacodylate buffer (pH 7.4) and then stored overnight in the fixative at 4°C. The samples were then washed with 0.1 M cacodylate buffer, treated for 1 hour with 1% (wt/vol) OsO4 at 4°C, washed with 0.1 M cacodylate buffer and Milli-Q water, and stained with 1% (wt/vol) uranyl acetate in Mili-Q water. After a new washing step, samples were dehydrated in increasing concentrations of ethanol (70%, 80%, 90%,100%), embedded in epoxy resin (LX-112, Ladd Research) and polymerized at 60°C. Sections (100 nm thick) were collected on mesh-100 copper EM grids covered with a carbon-coated Pioloform layer and post-stained with 7% (wt/vol) uranyl acetate and Reynold’s lead citrate. The samples were examined in a Twin transmission electron microscope (Thermo Fisher Scientific (formerly FEI)) operated at 120 kV and images were collected with a OneView 4k high-frame rate CMOS camera (Gatan).

### Compounds and antiviral screening assay

A 10-mM stock of Remdesivir (HY-104077; MedChemexpress) was dissolved in DMSO and stored in aliquots for single use at −80°C. Alisporivir was kindly provided by DebioPharm (Dr. Grégoire Vuagniaux, Lausanne, Switzerland; (44)) and a 20-mM stock was dissolved in 96% ethanol and stored in aliquots for single use at −20°C. A 20-mM chloroquine stock (C6628; Sigma) was dissolved in PBS and stored in aliquots for single use at −20°C. Pegylated interferon alpha-2a (PEG-IFN-α; Pegasys, 90 mcg, Roche) was aliquoted and stored at room temperature until further use. Vero E6 cells were seeded in a 96-well flat bottom plates in 100 µl at a density of 10,000 cells/well and grown overnight at 37°C. Two-fold serial dilutions of compounds were prepared in EMEM with 2% FCS and 50 µl was added to the cells 30 min prior to infection. Subsequently, half of the wells were infected with 300 PFU each of SARS-CoV or SARS-CoV-2 in order to evaluate inhibition of infection, while the other wells were used to in parallel monitor the (potential) cytotoxicity of compounds. Each compound concentration was tested in quadruplicate and each assay plate contained the following controls: no cells (background control), cells only treated with medium (mock infection for normalization), infected/untreated cells and infected/solvent-treated cells (infection control). At 3 days p.i., 20 μL/well of CellTiter 96 Aqueous Non-Radioactive Cell Proliferation reagent (Promega) was added and plates were incubated for 2 h at 37°C. Reactions were stopped and virus inactivated by adding 30 µl of 37% formaldehyde. Absorbance was measured using a monochromatic filter in a multimode plate reader (Envision; Perkin Elmer). Data was normalized to the mock-infected control, after which EC_50_ and CC_50_ values were calculated with Graph-Pad Prism 7.

## RESULTS

### Rapid adaptation of SARS-CoV-2 BetaCoV/Australia/VIC01/2020 during passaging in Vero E6 cells

SARS-CoV-2 isolate BetaCoV/Australia/VIC01/2020 was received as a stock derived from two consecutive passages in Vero/hSLAM cells (34). The virus was then propagated two more times at low MOI in Vero E6 cells, in which it caused a severe cytopathic effect (CPE). We also attempted propagation in HuH7 cells, using the same amount of virus or a ten-fold larger inoculum, but did not observe any cytopathology after 72 h (data not shown). At 24 h p.i., immunofluorescence microscopy (see below) revealed infection of only a small percentage of the HuH7 cells, without any clear spread to other cells occurring in the next 48 h. We therefore conclude that infection of HuH7 cells does not lead to a productive SARS-CoV-2 infection and deemed this cell line unsuitable for further SARS-CoV-2 studies.

The infectivity titre of the Leiden-p2 stock grown in Vero E6 cells was analysed by plaque assay, after which we noticed a mixed plaque phenotype (∼1:3 ratio of small versus large (plaques; data not shown) while a virus titre of 7 × 10^6^ PFU/ml was calculated. To verify the identity and genome sequence of the SARS-CoV-2/p2 virus stock, we isolated genomic RNA from culture supernatant and applied next-generation sequencing (NGS; see methods for details). The resulting consensus sequence was found to be identical to the sequence previously deposited in GenBank (accession number MT007544.1) (34), with one exception (see below). Compared to the SARS-CoV-2 GenBank reference sequence (NC_045512.3) (38) and other field isolates (29), isolate BetaCoV/Australia/VIC01/2020 exhibits >99.9% sequence identity. In addition to synonymous mutations in the nsp14-coding sequence (U19065 to C) and S protein gene (U22303 to G), ORF3a contains a single non-synonymous mutation (G26144 to U). Strikingly, the 3’ untranslated region (UTR) contains a 10-nt deletion (nt 29750-29759; CGAUCGAGUG) located 120 nt upstream of the genomic 3’ end, which is not present in other SARS-CoV-2 isolates thus far (>670 SARS-CoV2 sequences present in GenBank on April 17, 2020).

In about 71% of the 95,173 p2 NGS reads covering this position, we noticed a G23607 to A mutation encoding an Arg682 to Gln substitution near the so-called S1/S2 cleavage site of the viral S protein (see Discussion), with the other 29% of the reads being wild-type sequence. As this ratio approximated the observed ratio between large and small plaques, we performed a plaque assay on the p1 virus stock (Fig. 1a, leftmost well) and picked multiple plaques of each size, which were passaged three times in Vero E6 cells while monitoring their plaque phenotype. Interestingly, for several of the small-plaque virus clones (like S5; Fig. 1a) we observed rapid conversion to a mixed or large-plaque phenotype during these three passages, while large-plaque virus clones (like L8) stably retained their plaque phenotype (Fig. 1a). NGS analysis of the genome of a large-plaque p1 virus (L8p1) revealed that >99% of the reads in the S1/S2 cleavage site region contained the G23607 to A mutation described above. No other mutations were detected in the genome, thus clearly linking the Arg682 to Gln substitution in the S protein to the large-plaque phenotype observed for the L8p1 virus.

Next, we also analysed the genomes of the p1, p2 and p3 viruses derived from a small-plaque (S5) that was picked. This virus clone retained its small-plaque phenotype during the first passage (Fig. 1a; S5p1), but began to yield an increasing proportion of large(r) plaques during subsequent passages. Sequencing of S5p2 (Fig 1b) revealed a variety of low-frequency reads with mutations near the S1/S2 cleavage site motif (aa 681-687; PRRAR↓SV), with G23607 to A (specifying the Arg682 to Gln substitution) again being the dominant one (in ∼0.9% of the reads covering nt 23,576 to 23,665 of the genome). At lower frequencies single-nucleotide changes specifying Arg682 to Trp and Arg683 to Leu substitutions were also detected. Furthermore, a 10-aa deletion (residues 679-688) that erases the S1/S2 cleavage site region was discovered, as well as a 5-aa deletion (residues 675-679) immediately preceding that region. The amount of large plaques increased substantially upon the next passage, with NGS revealing the prominent emergence of the mutants containing the 10-aa deletion or the Arg682 to Gln point mutation (∼22% and ∼12% of the reads, respectively), and yet other minor variants with mutations in the PRRAR↓SV sequence being discovered. Taken together these data clearly link the large-plaque phenotype of SARS-CoV-2 to the acquisition of mutations in this particular region of the S protein, which apparently provides a strong selective advantage during passaging in Vero E6 cells.

### Comparative kinetics of SARS-CoV and SARS-CoV-2 replication in Vero E6 cells

To our knowledge, a detailed comparison of SARS-CoV-2 and SARS-CoV replication kinetics in cell culture has not been reported so far. Therefore, we infected Vero E6 cells with the SASR-CoV-2/p2 virus stock at high m.o.i. to analyse viral RNA synthesis, protein expression and the release of infectious viral progeny (Fig. 2a). This experiment was performed using 4 replicates per time point and for comparison we included the SARS-CoV Frankfurt-1 isolate (Drosten, Gunther et al. 2003), which has been used in our laboratory since 2003. During the early stages of infection (until 8 h p.i.), the growth curves of the two viruses were similar, but subsequently cells infected with SARS-CoV clearly produced more infectious progeny (about 50-fold more) than SARS-CoV-2-infected cells, with both viruses reaching their plateau by about 14 h p.i. As shown in Fig. 2b, despite its transition to a mainly large-plaque phenotype, the largest SARS-CoV-2/p3 plaques were still substantially smaller than those obtained with SARS-CoV Frankfurt-1.

**Fig. 2.**
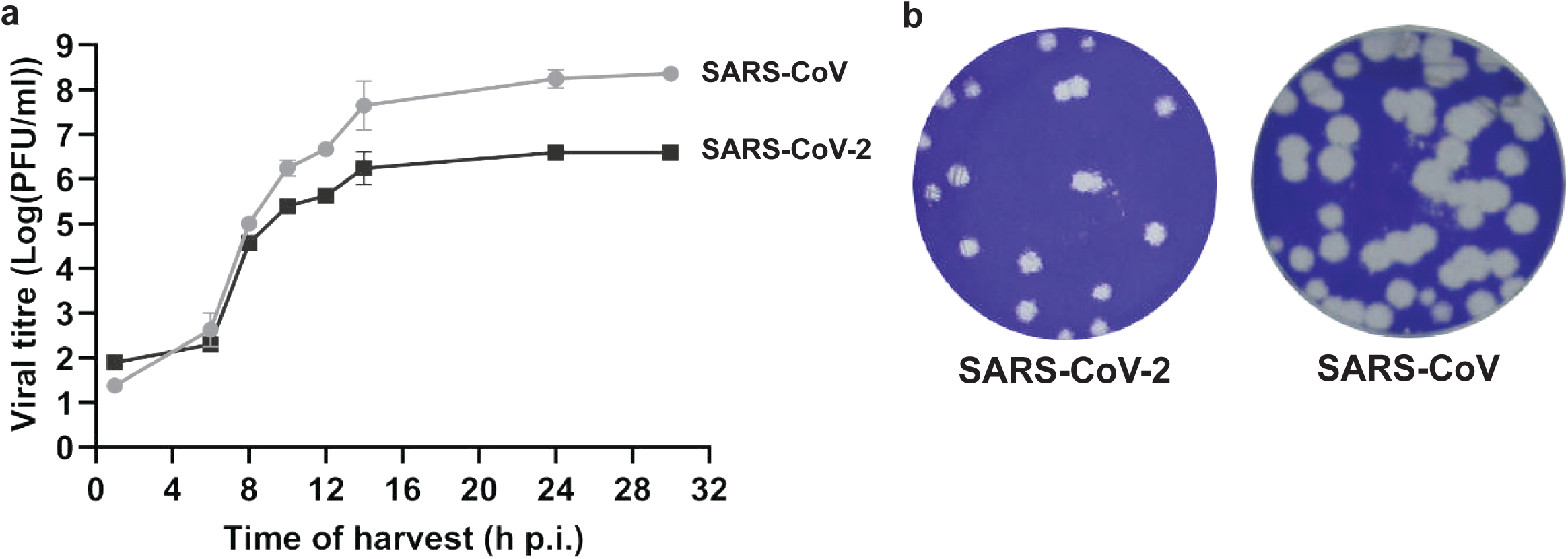
Comparison of SARS-CoV-2 and SARS-CoV replication kinetics in Vero E6 cells. (a) Growth curve showing the release of infectious viral progeny into the medium of infected Vero E6 cells (m.o.i. 3), as determined by plaque assay (n = 4; mean ± sd is presented). (b). Comparison of SARS-CoV-2 Australia/VIC01/2020 and SARS-CoV Frankfurt-1 plaque phenotype in Vero E6 cells.

In parallel, we analysed the kinetics of viral RNA synthesis by isolating intracellular viral RNA, subjecting it to agarose gel electrophoresis and visualizing the various viral mRNA species by in-gel hybridization with a ^32^P-labeled oligonucleotide probe recognizing a fully conserved 19-nt sequence located 30 nt upstream of the 3’ end of both viral genomes (Fig. 3a). This revealed the anticipated presence of the genomic RNA and eight subgenomic mRNAs, together forming the well-known 5’- and 3’-coterminal nested set of transcripts required for full CoV genome expression.

**Fig. 3.**
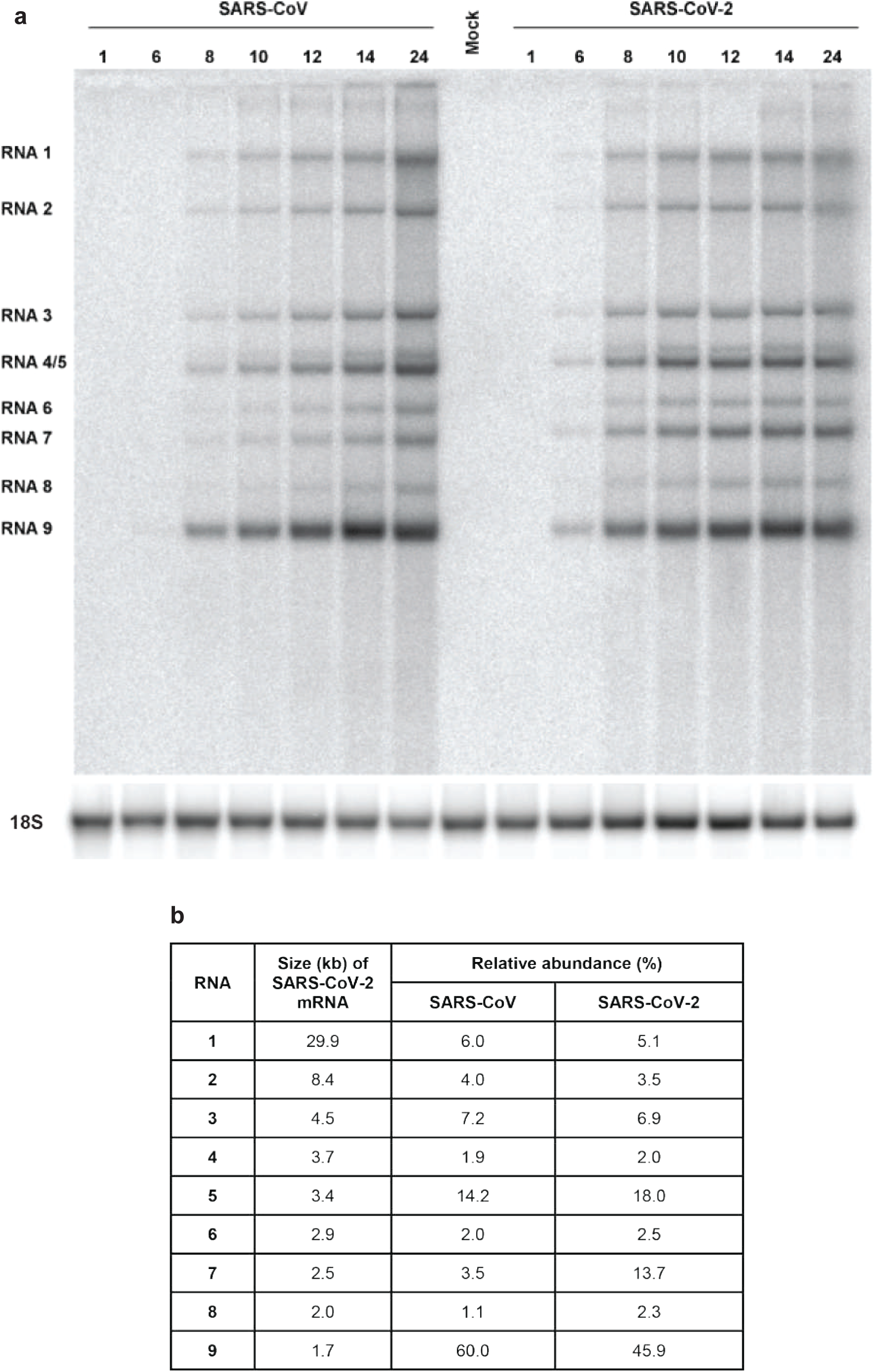
Kinetics of SARS-CoV-2 and SARS-CoV RNA synthesis in infected Vero E6 cells. (a) Hybridization analysis of viral mRNAs isolated from SARS-CoV-2- and SARS-CoV-infected Vero E6 cells, separated in an agarose gel and probed with a radiolabelled oligonucleotide recognizing the genome and subgenomic mRNAs of both viruses. Subsequently, the gel was re-hybridized to a probe specific for 18S ribosomal RNA, which was used as a loading control. (b) Analysis of the relative abundance of each of the SARS-CoV-2 and SARS-CoV transcripts. Phosphorimager quantification was performed for the bands of the samples isolated at 12, 14 and 24 h p.i., which yielded essentially identical relative abundances. The table shows the average of these three measurements. SARS-CoV-2 mRNA sizes were calculated on the basis of the position of the leader and body transcription-regulatory sequences (ACGAAC) in the viral genome (Sawicki and Sawicki 1995, Xu, Hu et al. 2003).

In general, for both viruses, the accumulation of viral RNAs followed the growth curves depicted in Fig. 2a. The relative abundance of the individual RNAs was determined using the 12, 14 and 24 h p.i. samples (averages presented in Fig. 3b) and found to be largely similar, with the exception of SARS-CoV-2 mRNAs 7 and 8, which accumulated to about 4 and 2 times higher levels, respectively. Strikingly, in spite of the ultimately lower yield of infectious viral progeny, SARS-CoV-2 RNA synthesis was detected earlier and reached an overall level exceeding that of SARS-CoV.

We also monitored viral protein production by Western blot analysis using antisera targeting a non-structural (nsp3) and structural (N) protein. As expected from the RNA analysis, the accumulation of both viral proteins increased with time, and was detected somewhat earlier for SARS-CoV-2 than for SARS-CoV (data not shown). Overall, we conclude that in Vero E6 cells, SARS-CoV-2 produces levels of intracellular RNA and proteins that are at least comparable to those of SARS-CoV, although this does not translate into the release of equal amounts of infectious viral progeny (Fig. 2a).

### Cross-reactivity of antisera previously raised against SARS-CoV targets

To be able to follow virus replication in SARS-CoV-2-infected cells more closely, we explored cross-reactivity of a variety of antisera previously raised against SARS-CoV targets, in particular a variety of nsps. In an earlier study, many of those were found to cross-react also with the corresponding MERS-CoV targets (35), despite the relatively large evolutionary distance between MERS-CoV and SARS-CoV. Based on the much closer relationship with SARS-CoV-2, similar or better cross-reactivity of these SARS-CoV reagents was expected, which was explored using immunofluorescence microscopy.

Indeed, most antisera recognizing SARS-CoV nsps that were tested (nsp3, nsp4, nsp5, nsp8, nsp9, nsp13, nsp15) strongly cross-reacted with the corresponding SARS-CoV-2 target (Fig. 4 and Table 1), the exception being a polyclonal nsp6 rabbit antiserum. Likewise, both a polyclonal rabbit antiserum and mouse monoclonal antibody recognizing the N protein cross-reacted strongly (Fig. 4b and Table 1). The same was true for a rabbit antiserum raised against a C-terminal peptide of the SARS-CoV M protein (Fig 4e). Labelling patterns were essentially identical to those previously documented for SARS-CoV (Stertz, Reichelt et al. 2007, Knoops, Kikkert et al. 2008), with nsps accumulating in the perinuclear region of infected cells, where the elaborate membrane structures of the viral ROs are formed (Fig. 4a, c, d). Punctate structures in the same area of the cell were labelled using an antibody recognizing double-stranded RNA (dsRNA), which presumably recognizes replicative intermediates of viral RNA synthesis (45, 46). The N protein signal was diffusely cytosolic (Fig. 4b), whereas the M protein labelling predominantly showed the expected localization to the Golgi complex (Fig. 4e), where the protein is known to accumulate (47).

**Fig. 4.**
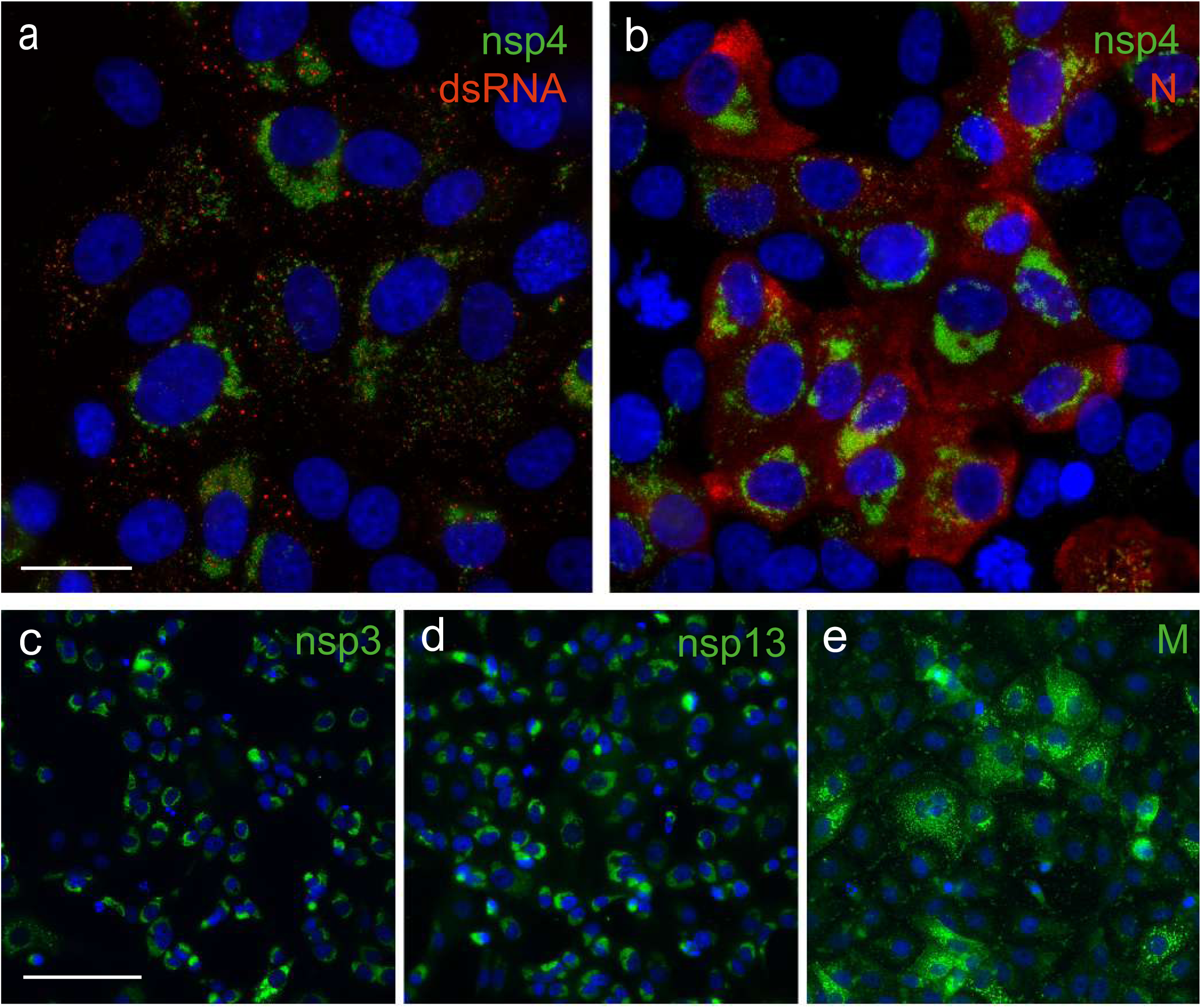
Cross-reactivity of antisera raised against SARS-CoV structural and non-structural proteins. Selected antisera previously raised against SARS-CoV nsps and structural proteins cross-react with corresponding SARS-CoV-2 proteins. SARS-CoV-2-infected Vero E6 cells (m.o.i. of 0.3) were fixed at 12 or 24 h p.i. For immunofluorescence microscopy, cells were (double)labelled with (a) a rabbit antiserum recognising nsp4 and a mouse mAb recognising dsRNA; (b) anti-nsp4 rabbit serum and a mouse mAb directed against the N protein; (c-e) rabbit antisera recognising against nsp3, nsp13 and the M protein, respectively. Nuclear DNA was stained with Hoechst 33258. Bar, 20 µm.

### Ultrastructural characterisation of SARS-CoV-2-infected cells

We next used electron microscopy to investigate the ultrastructural changes that SARS-CoV-2 induces in infected cells, and focused on the membranous replication organelles (ROs) that supports viral RNA synthesis and on the assembly and release of new virions (Fig. 5). Compared to mock-infected control cells (Fig. 5a-b), various distinct membrane alterations were observed in cells infected with either SARS-CoV or SARS-CoV-2 (Fig. 5c-j). At 6 h p.i., larger regions with membrane alterations were found particularly in cells infected with SARS-CoV-2 (data not shown), which may align with the somewhat faster onset of intracellular RNA synthesis in SARS-CoV2-infected Vero E6 cells (Fig. 3a). From 8 h p.i onwards, SARS-CoV- and SARS-CoV-2-infected cells appeared more similar (Fig. 5c-f and 5g-j). Double-membrane vesicles (DMVs) were the most prominent membrane alteration up to this stage (Fig. 5d-e and and 5h-i, asterisks). In addition, convoluted membranes (45) were readily detected in SARS-CoV-infected cells, while zippered ER (25, 48, 49) appeared to be the predominant structure in SARS-CoV-2-infected cells (Fig. 5e and 5i, white arrowheads). As previously described for SARS-CoV (45), also SARS-CoV-2-induced DMV appeared to fuse through their outer membrane, giving rise to vesicles packets that increased in numbers as infection progressed (Fig 5f and 5k, white asterisks). Virus budding near the Golgi apparatus, presumably into smooth membranes of the ER-Golgi intermediate compartment (ERGIC) (50-52), was frequently observed at 8 h p.i. (Fig. 5k-l and 5o-p). This step is followed by transport to the plasma membrane and release of virus particles into extracellular space. By 10 h p.i., released progeny virions were abundantly detected around all infected cells (Fig. 5m-n and 5q-r). Interestingly, whereas spikes were clearly present on SARS-CoV progeny virions, a relatively large proportion of SARS-CoV-2 particles seemed to carry few or no visible spike projections on their surface, perhaps suggesting a relatively inefficient incorporation of spike proteins into SARS-CoV-2 virions. This could potentially reduce the yield of infectious particles and may contribute to the lower progeny titres obtained for this virus (Fig. 2a).

**Fig. 5.**
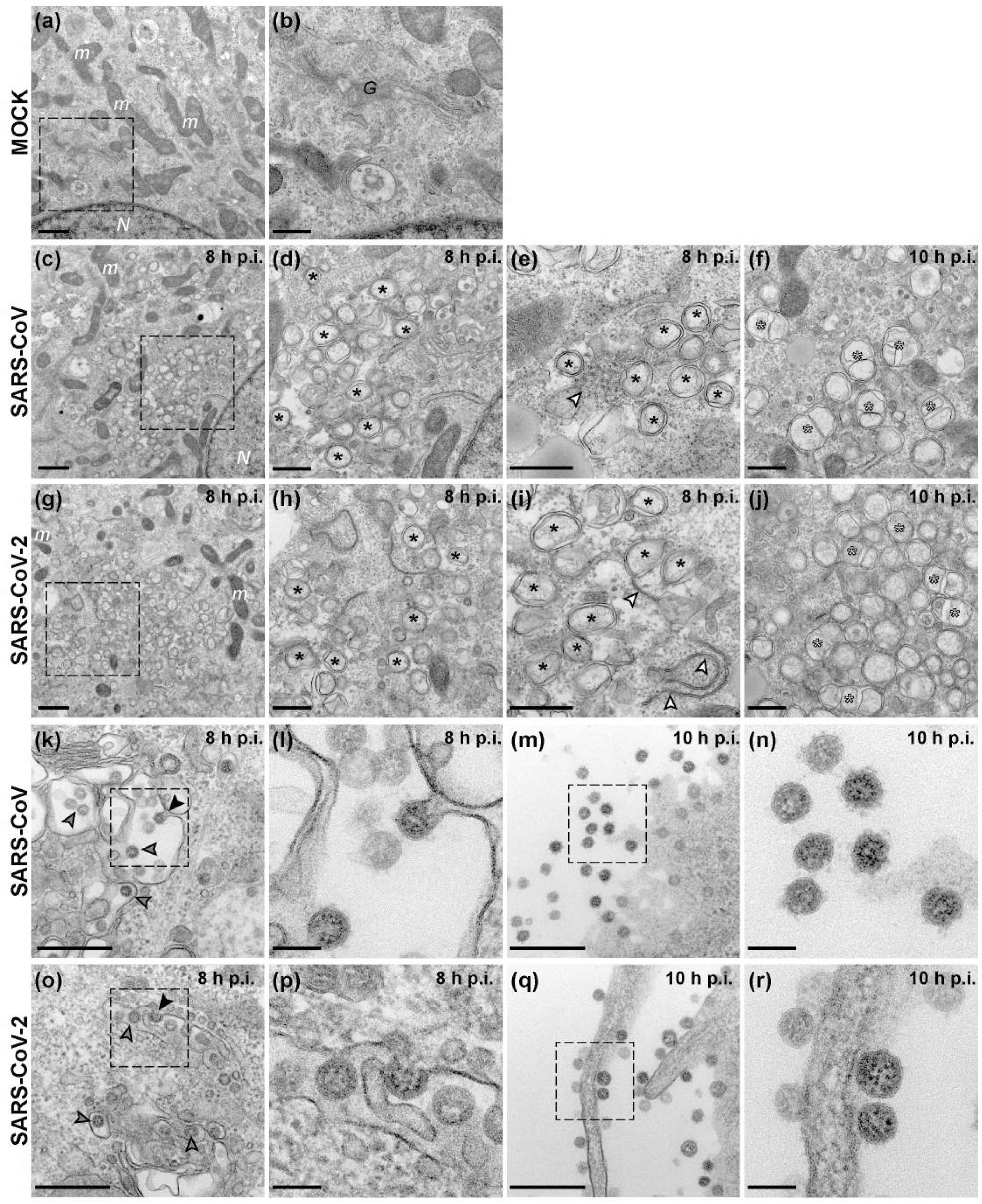
Visualisation of SARS-CoV-2 and SARS-CoV infection by electron microscopy. Electron micrographs of Vero E6 cells infected with either SARS-CoV-2 or SARS-CoV at the indicated time points (c-r). Images from a mock-infected cell are included for comparison (a-b). (c-j) RRegions containing viral replication organelles. These virus-induced structures accumulated in large clusters in the perinuclear region by 8 h p.i. (c, g, boxed regions enlarged in d and h, respectively). These regions primarily contained DMVs (d-e, h-i, black asterisks). Additionally, virus-induced convoluted membranes (e, white arrowhead) were observed in SARS-CoV infection, whereas zippered ER (i, white arrowheads) appeared to be more common in SARS-CoV-2-infected cells. At 10 h p.i., vesicle packets (f, j, white asterisks), which seem to arise by fusion of two or more DMVs through their outer membrane, became abundant in the RO regions. (k-r) Examples of virion assembly and release in infected cells. Virus particles budding into membranes of the ERGIC (k-l, o-p, arrowheads). The black arrowheads in the boxed areas highlight captured budding events, enlarged in l and p. Subsequently, virus particles are transported to the plasma membrane which, at 10 h p.i., is surrounded by a large number of released virions (m, q, boxed areas enlarged in n and r, respectively). N, nucleus; m, mitochondria; G, Golgi apparatus. Scale bars: 1 µm (a, c, g); 500 nm (b, d-f, h-j, k, m, o, q); 100 nm (l, n, p, r).

### Establishing a CPE-based assay to screen compounds for anti-SARS-CoV-2 activity

In order to establish and validate a CPE-based assay to identify potential inhibitors of SARS-CoV-2 replication, we selected four previously identified inhibitors of CoV replication: Remdesivir (53, 54), chloroquine (55, 56), Alisporivir (57, 58) and pegylated interferon alpha (PEG-IFN-α) (35, 59). Cells were infected at low MOI to allow for multiple cycles of replication. After three days, a colorimetric cell viability assay (60) was used to measure drug toxicity and inhibition of virus replication in mock- and virus-infected cells, respectively. With the exception of PEG-IFN-α, the inhibition of virus replication by compounds tested and the calculated half-maximal effective concentrations (EC_50_) were similar for SARS-CoV and SARS-CoV-2. For Remdesivir, we obtained higher EC_50_ values for SARS-CoV-2 and SARS-CoV (4.4 and 4.5 µM, respectively; Fig. 6a) than previously reported by others, but this may be explained by technical differences like a longer assay incubation time (72 h instead of 48 h) and the use of a different read-out (cell viability instead of qRT-PCR or viral load). Based on the obtained CC_50_ values of >100 µM, a selectivity index >22.5 was calculated. Chloroquine potently blocked virus infection at low-micromolar concentrations, with an EC_50_ value of 2.3 µM for both viruses (CC_50_ >100 µM, SI >45.5; Fig. 6b). Alisporivir, a known inhibitor of different groups of RNA viruses, was previously found to effectively reduce the production of CoV progeny. In this study, we measured EC_50_ values of 4.9 and 4.3 µM for SARS-CoV-2 and SARS-CoV, respectively (Fig. 6c; CC_50_>100 µM, SI >20). Treatment with PEG-IFN-α completely inhibited replication of SARS-CoV-2, even at the lowest dose of 7.8 ng/ml (Fig. 6d). In line with previous results (35, 59), SARS-CoV was much less sensitive to PEG-IFN-α treatment, yielding only partial inhibition at all concentrations tested (from 7.8 to 1000 ng/ml). Overall, we conclude that Vero E6 cells provide a suitable basis to perform antiviral compound screening and select the most promising hits for in-depth mechanistic studies and further development.

**Fig. 6.**
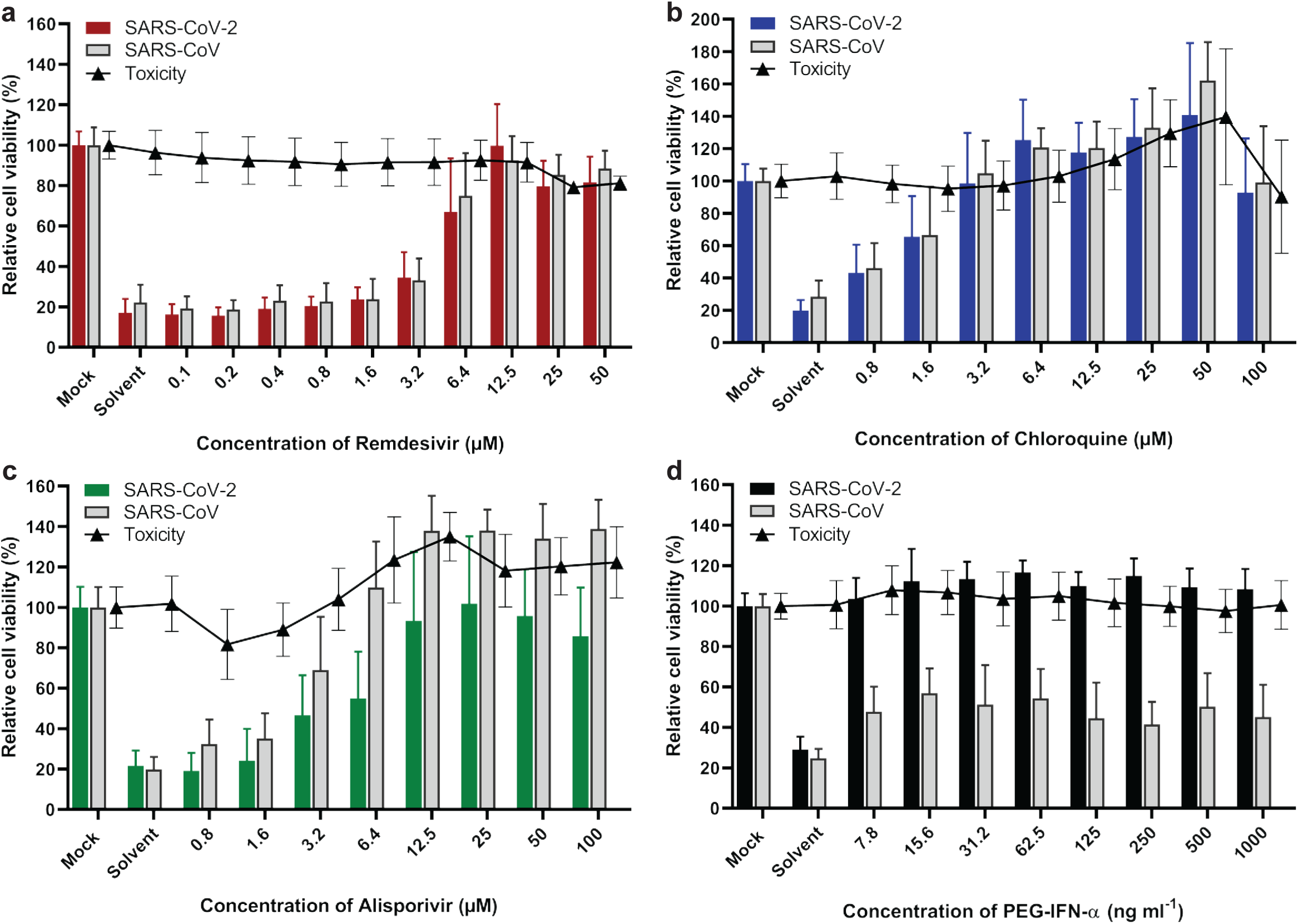
Assay to screen for compounds that inhibit SARS-CoV-2 replication. Inhibition of SARS-CoV-2 replication (coloured bars) was tested in Vero E6 cells by developing a CPE-reduction assay and evaluating several previously identified inhibitors of SARS-CoV, which was included for comparison (grey bars). For each compound a two-fold serial dilution series in the low-micromolar range was tested; (a) Remdesivir, (b) chloroquine, (c) Alisporivir and (d) pegylated interferon alpha-2. Cell viability was assayed using the CellTiter 96® Aqueous One Solution cell proliferation assay (MTS assay). Compound toxicity (solid line) was evaluated in parallel using mock-infected, compound-treated cells. The graphs show the results of 3 independent experiments, each performed using quadruplicate samples (mean ± SD are shown).

## Discussion

In this report, we describe a comparative analysis of the replication features of SARS-CoV-2 and SARS-CoV in Vero E6 cells, one of the most commonly used cell lines for studying these two viruses. However, in contrast to the stable phenotype exhibited by SARS-CoV during our 17 years of working with this virus in these cells, SARS-CoV-2 began to exhibit remarkable phenotypic variation in plaque assays within a few passages after its isolation from clinical samples (Fig. 1a). In addition to the BetaCoV/Australia/VIC01/2020 isolate used in this study, similar observations were made for a variety of other clinical isolates (data not shown). To establish the genetic basis for the observed plaque size heterogeneity, small and large plaques were picked and the resulting virus clones were passaged repeatedly and analysed using NGS. The consensus sequences obtained for S5p1 and L8p1, which differed by a single nucleotide substitution in the S protein gene, clearly established that a single S protein mutation (Arg682 to Gln) was responsible for the observed plaque size difference. This mutation is localized near the so-called ‘furin-like’ S1/S2 cleavage site (Fig. 1b) (61) in the S protein (62). This sequence constitutes a (potential) processing site that is present in a subset of CoVs (including SARS-CoV-2 and MERS-CoV) but is lacking in others, like SARS-CoV and certain bat CoVs (61, 63). This polybasic motif (PRRAR↓SV, in SARS-CoV-2) can be recognized by intracellular furin-like proteases during viral egress and its cleavage is thought to prime the S protein for fusion and entry (64), which also requires a second cleavage event to occur at the downstream S2’ cleavage site (61). In general, the presence of the furin-like cleavage site does not appear to be critical for successful CoV infection. Using pseudotyped virions carrying mutant S proteins of SARS-CoV (65) or SARS-CoV-2 (66), it was shown that its presence minimally impacts S protein functionality. In the SARS-CoV S protein, an adjacent sequence that is conserved across CoVs can be cleaved by other host proteases like cathepsin L or TMPRSS2 (67-69), thus providing an alternative pathway to trigger viral entry. Possibly, this pathway is also employed by our Vero E6-cell adapted SARS-CoV-2 mutants that have lost the furin-like cleavage site, like clone L8p1 and multiple variants encountered in S5p3 (Fig. 1a). These variants contain either single point mutations or deletions of 5 to 10 aa (Fig. 1b), resembling variants recently reported by other laboratories (30, 70, 71). Interestingly similar changes were also observed in some clinical SARS-CoV-2 isolates that had not been passaged in cell culture (70). It is currently being investigated why mutations that inactivate the furin-like cleavage site provide such a major selective advantage during SARS-CoV-2 passaging in Vero E6 cells and how this translates into the striking large-plaque phenotype documented in this paper.

An additional remarkable feature confirmed by our re-sequencing of the BetaCoV/Australia/VIC01/2020 isolate of SARS-CoV-2 is the presence of a 10-nt deletion in the 3’ UTR of the genome (34). Screening of other available SARS-CoV-2 genome sequences indicated that the presence of this deletion apparently is unique for this particular isolate, and likely represents an additional adaptation acquired during cell culture passaging. This deletion maps to a previously described “hypervariable region” in the otherwise conserved 3’ UTR, and in particular to the so-called s2m motif (72) that is conserved among CoVs and also found in several other virus groups (73, 74). The s2m element has been implicated in the binding of host factors to viral RNAs, but its exact function has remained enigmatic thus far. Strikingly, for the mouse hepatitis coronavirus the entire hypervariable region (including s2m) was found to be dispensable for replication in cell culture, but highly relevant for viral pathogenesis in mice (72). Although the impact of this deletion for SARS-CoV-2 remains to be studied in more detail, these previous data suggest that this mutation need not have a major impact on SARS-CoV-2 replication in Vero E6 cells. This notion is also supported by the fact that the results of our antiviral screening assays (Fig. 6) correlate well with similar studies performed with other SARS-CoV-2 isolates (54, 75, 76). Clearly, this could be different for *in vivo* studies, for which it would probably be better to rely on SARS-CoV-2 isolates not carrying this deletion in their 3’ UTR.

Vero E6 cells are commonly used to isolate, propagate, and study SARS-CoV-like viruses as they support viral replication to high titres (77-81). This may be due to a high expression level of the ACE-2 receptor (82) that is used by both SARS-CoV-2 and SARS-CoV (9) and/or the fact that they lack the ability to produce interferon (83, 84). It will be interesting to evaluate whether there is a similarly strong selection pressure to adapt the S1/S2 region of the S protein when SARS-CoV-2 is passaged in other cell types. Such studies are currently in progress in our laboratory and already established that HuH7 cells may be a poor choice, despite the fact that they were used for virus propagation (9, 85) and antiviral screening in other studies (54, 86). Immunolabelling of infected HuH7 cells (data not shown) revealed non-productive infection of only a small fraction of the cells and a general lack of cytopathology. While other cell lines are being evaluated, as illustrated above, the monitoring of the plaque phenotype (plaque size and homogeneity) may provide a quick and convenient method to assess the composition of SARS-CoV-2 stocks propagated in Vero E6 cells, at least where it concerns the evolution of the S1/S2 region of the S protein.

Given the ongoing SARS-CoV-2 pandemic, the detailed characterization of its replication cycle is an important step in understanding the molecular biology of the virus and defining potential targets for inhibitors of replication. The cross-reacting antisera described in this study (Table 1) will be a useful tool during such studies. In general, the subcellular localization of viral nsps and structural proteins (Fig. 4) and the ultrastructural changes associated with RO formation (Fig. 5) were very similar for the two viruses. We also observed comparable replication kinetics for SARS-CoV-2 and SARS-CoV in Vero E6 cells, although clearly lower final infectivity titres were measured for SARS-CoV-2 (∼50-fold lower; Fig. 2). Nevertheless, RNA synthesis could be detected somewhat earlier for SARS-CoV-2 and the overall amount of viral RNA produced exceeded that produced by SARS-CoV (Fig. 3). This may be indicative of certain assembly or maturation problems or of virus-host interactions that are different in the case SARS-CoV-2. These possibilities merit further investigation, in particular since our preliminary EM studies suggested intriguing differences with SARS-CoV where it concerns the presence of spikes on the surface of freshly released SARS-CoV-2 particles (Fig. 5n and 5r).

Our analysis of SARS-CoV-2 subgenomic mRNA synthesis revealed the increased relative abundance of mRNAs 7 and 8 (∼4- and ∼2-fold, respectively) when SARS-CoV-2 was compared to SARS-CoV. Mechanistically, these differences do not appear to be caused by extended base pairing possibilities of the transcription regulatory sequences that direct the synthesis of these two mRNAs (24). As in SARS-CoV, mRNA7 of SARS-CoV-2 encodes for two proteins, the ORF7a and ORF7b proteins, with the latter presumably being expressed following leaky ribosomal scanning (32). Upon its ectopic expression, the ORF7a protein has been reported to induce apoptosis via a caspase-dependent pathway (87) and/or to be involved in cell cycle arrest (88). The ORF7b product is a poorly studied integral membrane protein that has (also) been detected in virions (89). When ORF7a/b or ORF7a were deleted from the SARS-CoV genome, there was a minimal impact on the kinetics of virus replication *in vitro* in different cell lines, including Vero cells, and *in vivo* using mice. In another study, however, partial deletion of SARS-CoV ORF7b was reported to provide a replicative advantage in CaCo-2 and HuH7 cells, but not in Vero cells (90).

The SARS-CoV ORF8 protein is membrane-associated and able to induce endoplasmic reticulum stress (91, 92), although it has not been characterised in great detail in the context of viral infection. Soon after the emergence of SARS-CoV in 2003, a conspicuous 29-nt (out-of-frame) deletion in ORF8 was noticed in late(r) human isolates, but not in early human isolates and SARS-like viruses obtained from animal sources (93-95). Consequently, loss of ORF8 function was postulated to reflect an adaptation to the human host. The re-engineering of an intact ORF8, using a reverse genetics system for the SARS-CoV Frankfurt-1 isolate, yielded a virus with strikingly enhanced (up to 23-fold) replication properties in multiple systems (96). Clearly, it remains to be established that the increased synthesis of mRNAs 7 and 8 is a general feature of SARS-CoV-2 isolates, and that this indeed also translates into higher expression levels of the accessory proteins encoded by ORFs 7a, 7b and 8. If confirmed, these differences definitely warrant an in-depth follow-up analysis as CoV accessory proteins in general have been shown to be important determinants of virulence. They may thus be relevant for our understanding of the wide spectrum of respiratory disease symptoms observed in COVID-19 patients (97).

Based on the close ancestral relationship between SARS-CoV-2 and SARS-CoV (98), one might expect that the patterns and modes of interaction with host antiviral defence mechanisms would be similar. However, our experiments with type I interferon treatment of Vero E6 cells (Fig. 6) revealed a clear difference, with SARS-CoV-2 being considerably more sensitive than SARS-CoV, as also observed by other laboratories (76). Essentially, SARS-CoV-2 replication could be inhibited by similarly low concentrations of PEG-IFN-alpha-2a that inhibit MERS-CoV replication in cell culture (35). Taken together, our data suggest that SARS-CoV-2 is less able to counter a primed type I IFN response than SARS-CoV (76, 99).

Previously identified inhibitors of CoV replication were used to further validate our cell-based assay for SARS-CoV-2 inhibitor screening. These compounds inhibited replication at similar low-micromolar concentrations and in a similar dose-dependent manner as observed for SARS-CoV (Fig. 6). Remdesivir is a prodrug of an adenosine analogue developed by Gilead Sciences. It was demonstrated to target the CoV RNA polymerase and act as a chain terminator (100-102). The clinical efficacy of Remdesivir is still being evaluated and, after some first encouraging results (103), worldwide compassionate use trials are now being conducted. Likewise, hydroxychloroquine and chloroquine have been labelled as potential “game changers” and are being evaluated for treatment of severe COVID-19 patients (104). Both compounds have been used to treat malaria and amebiasis (105), until drug-resistant *Plasmodium* strains emerged (106). These compounds can be incorporated into endosomes and lysosomes, raising the pH inside these intracellular compartments, which in turn may lead to defects in protein degradation and intracellular trafficking (68, 107). An alternative hypothesis to explain their anti-SARS-CoV activity is based on their impact on glycosylation of the ACE2 receptor that is used by SARS-CoV (56). Finally, as expected, the non-immunosuppressive cyclosporin A analogue Alisporivir inhibited SARS-CoV-2 replication, as demonstrated previously for SARS-CoV and MERS-CoV (58). Although the exact mode of action of this inhibitor it is unclear, it is thought to modulate CoV interactions with members of the cyclophilin family (108). Unfortunately, all of these *in vitro* antiviral activities should probably be classified as modest, emphasizing the urgency of large-scale drug repurposing and discovery programmes that target SARS-CoV-2 and coronaviruses at large.

## Abbreviations

SARS-CoV: severe acute respiratory syndrome coronavirus
CoV: Coronavirus
CPE: cytopathic effect
HCoV: human coronavirus
MERS-CoV: Middle East respiratory syndrome coronavirus;
nsp: non-structural protein
S protein: spike protein
ACE2: angiotensin-converting enzyme 2
NGS: next-generation sequencing
RO: replication organelle
DMV: Double-membrane vesicle
PEG-IFN-α: pegylated interferon alpha
UTR: untranslated region.

## Authors and contributors

NO, JD, MK, MB, IS and ES conceptualised the study. NO, TD, JZ, RL, YM and LC performed experimental work and contributed to analysis of the results and preparation of figures. NO, LC, JD, JV, IS and ES performed NGS and were involved in the bioinformatics analysis of the data. NO and ES wrote the manuscript, with input from all authors.

## Conflicts of interest

The authors declare that there are no conflicts of interest.

## Funding information

None.

## Acknowledgements

We thank various GenomeScan staff members for the pleasant and swift collaboration that facilitated the NGS and data analysis of the first SARS-CoV-2 samples. We are grateful to all members of the sections Research and Clinical Microbiology of the LUMC Department of Medical Microbiology for their collaborative support and dedication during the current pandemic situation. In particular, we thank Linda Boomaars, Peter Bredenbeek, Ien Dobbelaar, Martijn van Hemert, Sebenzile Myeni, Tessa Nelemans, Esther Quakkelaar, Ali Tas, Tessa Nelemans, Sjaak Voorden and Gijsbert van Willigen for their technical or administrative support, constructive discussions and/or scientific input.

## Notes

### Competing Interest Statement

The authors have declared no competing interest.

